# New avenues for potentially seeking microbial responses to climate change beneath Antarctic ice shelves

**DOI:** 10.1101/2023.12.13.571508

**Authors:** Aitana Llorenç Vicedo, Monica Lluesma Gomez, Ole Zeising, Thomas Kleiner, Johannes Freitag, Francisco J. Martínez-Hernández, Frank Wilhelms, Manuel Martínez-García

## Abstract

The signs of climate change are undeniable, and the impact of these changes on ecosystem function heavily depends on the response of microbes that underpin the food web. Antarctic ice shelf is a massive mass of floating ice that extends from the continent into the ocean, exerting a profound influence on global carbon cycles. Beneath Antarctic ice shelves, marine ice stores valuable genetic information, where marine microbial communities before the industrial revolution are archived. Here, in this proof-of-concept, by employing a combination of single-cell genomics and metagenomics, we have been able to sequence frozen microbial DNA (≍300 years old) stored in the marine ice core B15 collected from the Filchnner-Ronne Ice Shelf. Metagenomic data indicated that *Proteobacteria* and *Thaumarchaeota* (e.g. *Nitrosopumilus spp*.) followed by *Actinobacteria* (e.g. Actinomarinales) were abundant. Remarkably, our data allow us to ‘travel to the past’ and calibrate genomic and genetic evolutionary changes for ecologically relevant microbes and functions, such as *Nitrosopumilus* spp., preserved in the marine ice (≍300 years old) with those collected recently in seawater under an ice shelf (year 2017). The evolutionary divergence for the ammonia monooxygenase gene *amoA* involved in chemolithoautotrophy was about 0.88 amino acid and 2.8 nucleotide substitution rate per 100 sites in a century, while the accumulated rate of genomic SNPs was 2,467 per 1 Mb of genome and 100 years. Whether these evolutionary changes remained constant over the last 300 years or accelerated during post-industrial periods remains an open question that will be further elucidated.

**Importance:** Several efforts have been undertaken to predict the response of microbes under climate change, mainly based on short-term microcosm experiments under forced conditions. A common concern is that manipulative experiments cannot properly simulate the response of microbes to climate change, which is a long-term evolutionary process. In this proof-of-concept study with a limited sample size, we demonstrate a novel approach yet to be fully explored in science for accessing genetic information from putative past marine microbes preserved under Antarctic Ice shelves before Industrial revolution. This potentially allow us estimating evolutionary changes as exemplified in our study. We advocate for gathering a more comprehensive Antarctic marine ice core datasets across various periods and sites. Such a dataset would enable the establishment of a robust baseline, facilitating a better assessment of the potential effects of climate change on key genetic signatures of microbes.

## Introduction

Microbes sustain all other life forms and understanding how climate change affects this ‘giant microbial engin’ is paramount to assess and forecast the health of the ecosystem ^1–4^. Since industrial revolution, microbes face a novel combination of environmental challenges and global-scale anthropogenic perturbations of the Earth’s carbon and nutrient cycles that modify nearly every chemical, physical and biological characteristics that modulates the marine microbial growth. This poses the question of how microbes, which represent the pillar and foundation of the ecosystems will be reshaped. The signs of climate change are evident in polar environments ^5^, which are the regions of the world experiencing climate change (i.e., global warming) at the steepest rate. Indeed, microbes in these habitats might serve as “biosensors” ^6–13^. Several efforts have been made to predict the response of microbes under climate change ^8,14–18^. Some of them use short-term microcosm or in situ mesocosm experiments, incubating seawater from polar environments in laboratories under forced conditions, such as temperature rise^8,14–17,19^. Some of these studies have held controversy, since a common concern is that manipulative experiments cannot properly simulate the response of microbes to climate change^5^, which is a long-term evolutionary process. Alternatively, based on accumulative time-series data, ecological models that build theoretical frameworks have been used to forecast the fate of microbes in marine ecosystems ^5,20–26^.

Here, our study represents a novel approach that directly retrieve biological and genetic meaningful information from microbes preserved in the marine ice beneath the Filchnner-Ronne Ice shelf, which represent microbes inhabiting the past ocean with potential to shed some light into responses of microbes facing climate change.

The examination of biological records plays a crucial role in the field of environmental sciences, enabling us to comprehend the responses of organisms to specific environmental disturbances. Remarkably, beneath the Antarctic Ice shelves that cover an area of 1.561 million square kilometers, marine ice stores a yet to be explored valuable frozen genetic records of marine microbes with potential to open a “*time window*” that allow us to read some of the changes that microbes already experienced in the past. Antarctic ice shelves comprise a top freshwater layer of meteoric snow and a bottom layer of marine ice (i.e. frozen seawater) that can be as thick as several hundred meters^27–29^. Recently, it has been described that a functional and diverse microbial community similar to those in open meso-and bathypelagic ocean reside in the seawater under the Antarctic Ross Ice Shelf ^30^. Nowadays, we know that marine ice is formed beneath the Antarctic ice shelves trapping particles and microbes, and reach a depth of several hundred meters ^29,31–33^. The micro-structure of that marine ice, such as that beneath the Filchner-Ronne Ice Shelf, is uneven and lacks gas bubble and the typical brine drainage channels with liquid water that is common in surface sea ice ^34,35^, where it has been described that microbes might thrive. Thus, the marine ice under the Antarctic ice shelves is clearly distinct from surface sea ice, and microbes are trapped and frozen without available liquid water and gas preventing microbial growth. This marine ice also experiences complex dynamics of melting and freezing that accrete or reduce the layer of marine ice under the meteoric ice ^36,37^. Since the first discovery of marine ice in the Filchnner-Ronne Ice shelf ^29,32^ from two ice cores (B13 and B15; 65-72 mm diameter) drilled in 1992 near the ice shelf’s front ^38^ (Fig 1A), these have been stored in the Alfred Wegener Institute (AWI). For a complete physico-chemical characterization (e.g. salinity, etc..) of these ice cores, please refer to the original publications^29,32^. These ice cores serve as valuable repositories archiving microbial genetic signatures preserved from a past ocean that could provide insights into the fate of microbes in the context of climate change. In this proof-of-concept pilot study, by combining single-cell technologies and metagenomics, we have been able to sequence the microbial DNA from marine ice samples prior industrial revolution (≍300 years old). Our approach therefore has the potential to open new avenues in Microbiology for accessing frozen microbial DNA trapped in marine ice formed under the Antarctic ice shelves in the past.

**Figure 1.**
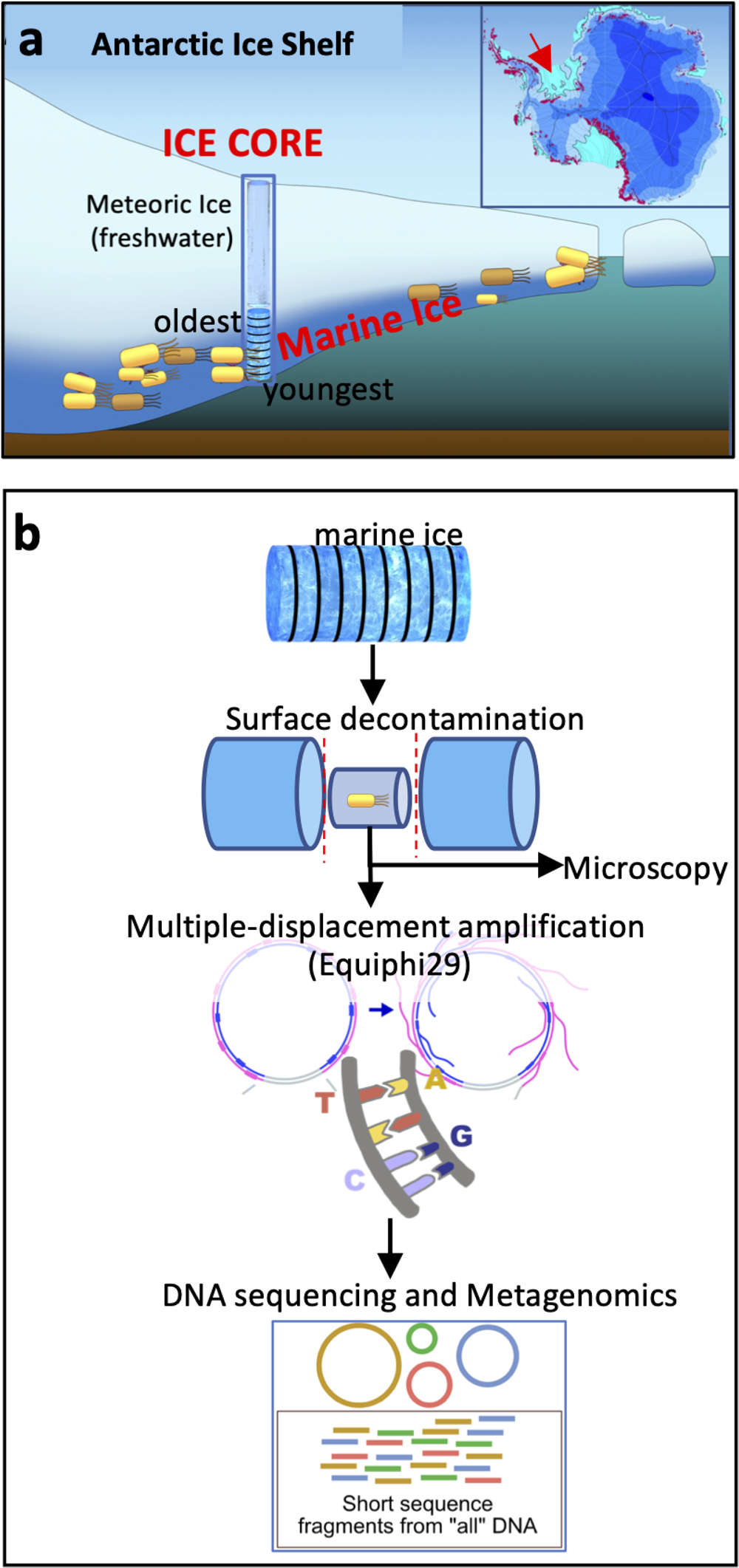
Analysis of microbiome preserved in marine ice beneath the Antarctic Ice Shelves. (a) Antarctic Ice Shelves (Light blue) are indicated in insert panel. Diagram showing the structure of an Antarctic ice shelf comprised of a meteoric ice layer (top) and marine ice formed at the bottom of the ice shelf. Youngest layers of marine ice are those in direct contact with seawater. Oldest marine ice is located at the interphase with meteoric ice. Red arrow in Antarctic map indicates the drilling site of ice core B15 used in this study (b) Schematic of the employed protocol that start with decontamination of the surface of the ice to obtain clean inner marine ice used for further experiments for molecular biology, confocal and electron microscopy.

## Results and discussion

### Decontamination of the Antarctic marine ice core B15

In this proof-of-concept pilot study, we have been able to sequence and explore the microbial genetic signatures stored in the marine ice beneath the Filchnner-Ronne Ice Shelf. For that, firstly we applied a previously reported robust protocol to decontaminate the surface of ice cores^9^ in sterile conditions that comprise 1) scraping the external part of the ice, 2) ethanol washing of the surface, and 3) sterile mQ water washing (Fig. 1B). This reported method proven to effectively decontaminate artificial ice cores contaminated with high concentrations of known bacteria (1X10^6^ cells) and viruses (4.48X10^7^ viruses) on its surface^9^. Furthermore, complementary methods used in single-cell and -virus genomic applications in our laboratory ^39–41^, which is the most extreme case scenario of ultra-low microbial biomass samples, were implemented to ensure no contamination with exogenous DNA. After decontamination, only the inner central part of the ice core (approximately 1 ml) was used for further microbial analysis as depicted in Fig. 1B and Fig. S1.

### Marine ice core sample dating and microscopy analyses

In order to estimate the age of the aggregated marine ice at the ice core B15, ice-flow model calculations were employed (see methods for a complete description). Briefly, we analyzed basal melt and accretion rates according to Adusumilli and colaborators^42^ along the flowline, accounting for the dynamic thinning from vertical strain (see ref ^43^ and Fig. S2-S4). Our data indicate that ice samples (≍135 m depth below meteoric ice) used in this pilot study from ice core B15 were approximately 275-328 years old before drilling (Fig. 2 and Figs. S2-S4). A similar methodology to estimate ice age based on ice-flow modelling has been extensively used in other studies on glaciology^44–47^.

**Figure 2.**
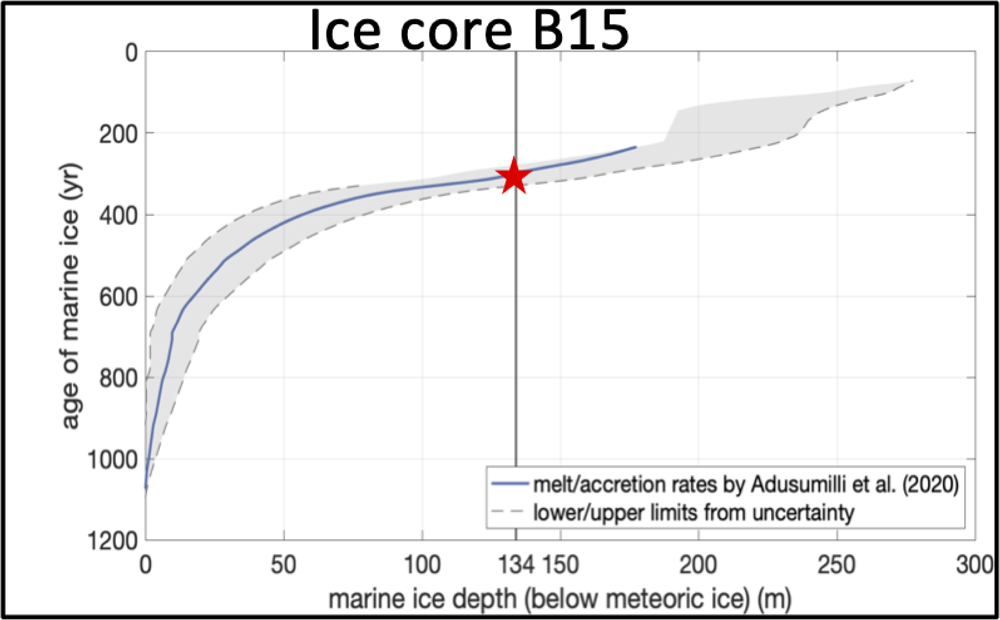
Dating based on ice-flow modeling of marine ice core B15 drilled in 1992 at the Filchner-Ronne Ice Shelf. Red star represents the marine ice samples used in our experiments (275-328 years old before drilling). Robust ice-flow model as explained in detail in methods and supplementary material were employed to estimate the age of the analyzed marine ice.

Next, we sought to explore the presence of intact microbial cell structures in the marine ice core B15 with common DNA fluorescent dye staining (SYBR Gold, DAPI and propidium iodide) and electron microscopy. For that, as explained above, we only used the inner central part of the marine ice after decontamination. Data from confocal and electron microscopy indicated that many of the observed cell-like structures were damaged and the detection of intact cell-like structures was a rare event (Fig. 3 and Figs. S5 and S6). In addition, particles trapped in marine ice were also abundant (Fig. S6). Unspecific fluorescence signal from dyes interacting with sediment particles has been commonly described^48,49^, which could explain here the abundant observed amorphous fluorescent stained structures (Fig. 3B). Likely the high pressure under hundred meters of ice layers exerts unfavorable preservation conditions of the cell structure, which overall preclude from a reliable cell counting.

**Figure 3.**
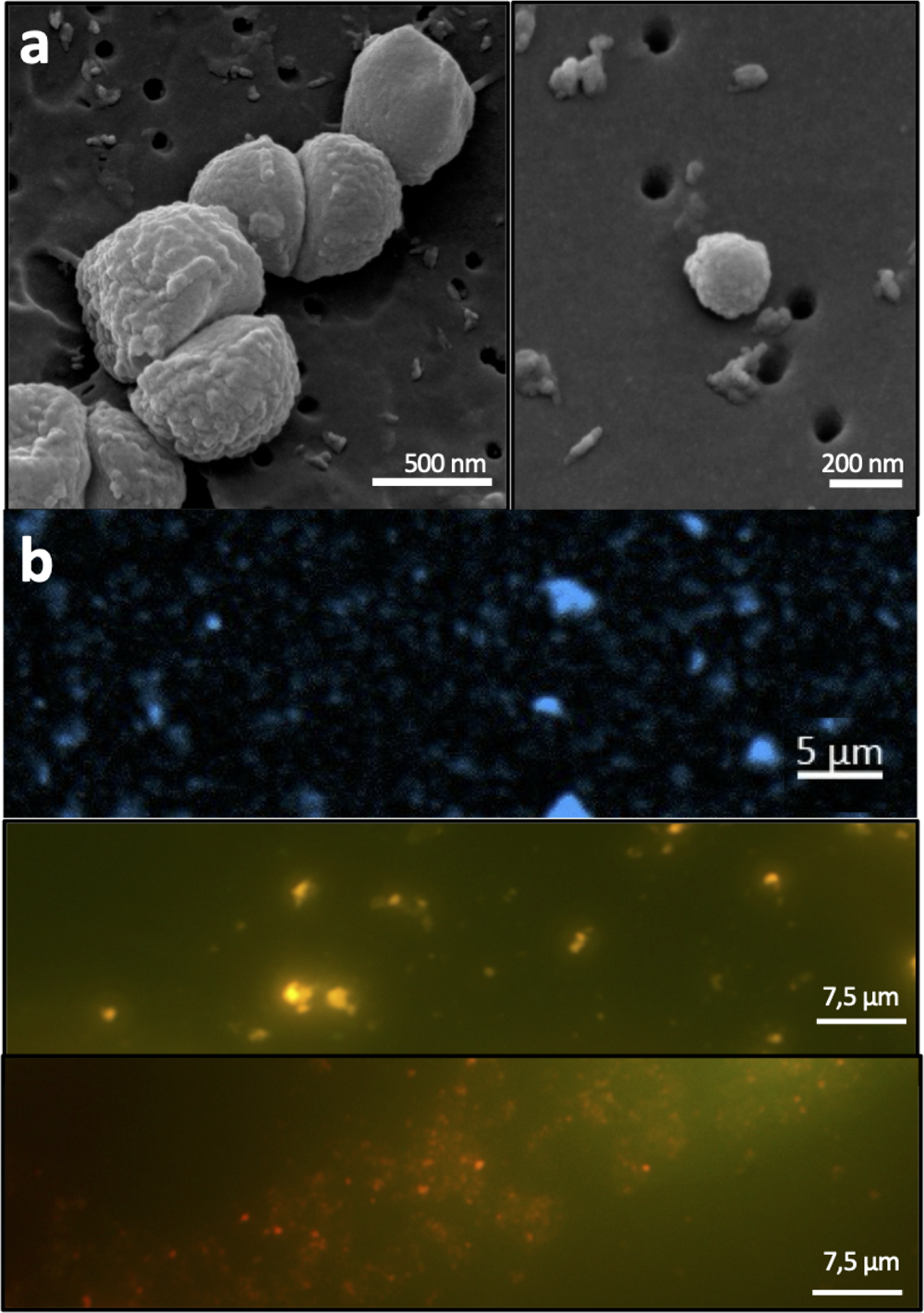
Microscopy analyses of the inner central part of the marine ice core B15. (a) Scanning electron microscopy (SEM) of marine ice. Observing nearly intact cell-like structures in the analyzed samples was very infrequent. Most of the observed objects were amorphous material with unclear cell-like structure. (b) In good agreement with SEM data showing high abundance of particles (see Supplementary Information), DNA dyes (DAPI, SYBR Gold and Propidium iodide) likely stain nonspecifically particles trapped in ice in addition to cell-like structures.

### Unveiling the genetic microbial traits stored in the marine ice core B15

Common DNA microbial extraction procedures and kits (see methods) recommended by gold standard procedures for microbiome analysis ^50,51^ failed to retrieve measurable DNA indicating that the amount of microbial DNA preserved in the sample was very limited, representing therefore an ultra-low biomass sample. In addition, it is important to remark that the available volume of marine ice from ice core B15 for experimentation was extremely low. To overcome these limitations, we applied single-cell genomic protocols to retrieve high molecular DNA present in ice (Fig. 1B and Figs. S7 and S8). For that, less than 1 ml of melted marine ice was used from different samples to perform real-time whole genome amplification and Illumina sequencing of the bulk microbial DNA allowing us to get access to the microbial metagenome preserved in the inner central part of the ice core (Fig. 1B). The fact that the enzyme phi29 successfully used during whole-genome amplification (i.e. multiple-displacement amplification) necessitates long DNA templates (a few kb) suggest that the DNA preserved at very low temperature in the analyzed samples was not short, degraded, nor fragmented. Data from two independent artificial blank ice cores of sterile mQ water built at AWI and University of Alicante and processed as the rest of samples demonstrated that potential exogenous DNA did not reach the inner central part of the ice core during the manipulation process (Fig. S9). This single-cell genomic method has been extensively proven to be robust for exploring the genetic information in biology and microbiology ^52^ despite tiny amount of DNA; even from a single cell or viral particle^40^.

According to 16S rRNA gene information from PCR screening of whole-genome amplified material (Fig. S8) and also from direct retrieval of 16S rRNA gene sequences^53^ from metagenomes (Fig. 4A) showed that marine microbes, such as *Actinomarinales* and *Nitrosopumilus spp,* and others (e.g. uncultured Gemmatinomonadota, uncultured Gammaproteobacteria and Chloroflexi), also detected in the seawater under the Ross Ice shelf ^30^, were present and preserved in the analyzed marine ice of 275-328 years old before drilling (Fig. 4). Metagenomic analysis of triplicate unassembled metagenomes showed that our methodology was robust and reproducible (MASH distance between metagenome replicates <0.05; values range from 0 to 1; 0-identical, 1-totally different). Overall metagenomic diversity of the microbiome preserved in the marine ice by means of Nonpareil diversity index was lower (value of ≍14.5) than that commonly measured in surface seawater/freshwater (values ≍20 or higher) or soils/sediments (values ≍25)^54^. Taxonomic assignment of unassembled metagenomic data from the different analyzed samples indicated that the preserved microbiome was dominated by *Proteobacteria* and *Thaumarchaeota* followed by marine *Actinobacteria* (mainly Actinomarinales) (Fig. 4B and Fig. S10). Remarkably, viral genetic information from Monoviridae, Caudoviricetes and Mimiviridae potentially infecting bacteria and also marine eukaryotes were also detected in our dataset (Fig. 4B). Data from gene annotation and metabolism (Fig. 5) indicate that genes involved on chemolithoautotrophy based on ammonia oxidation were present (e.g. *amoA, amoC, amoX* genes of *Nitrosopumilus spp.*); similarly to the microbiome described inhabiting in the dark ocean cavity under the Ross Ice Shelf^30^. Hydrogenases potentially involved in H_2_ oxidation were also detected. As expected, since light is not an available energy source for the microbiome thriving under hundreds of meters of ice, and in good agreement with data from beneath the Ross Ice Shelf, genes involved in oxygenic photosynthesis were not detected. Metabolism based on the oxidation of 1C molecules, as in the microbiome beneath the Ross Ice Shelf^30^, was predominant. Genes involved in hydrocarbon degradation were found as well. Rhodanases involved in sulfur transportation for assimilatory and dissimilatory sulphate reduction were also substantially detected in the metagenome. It is important to bear in mind that the microbial DNA recovered in the analyzed ice core samples of this study would originate from microbes that were inhabiting the ocean cavity beneath the Filchner-Ronne Ice Shelf, and were later trapped and frozen. Thus, all the above mentioned metabolic features would correspond to the metabolic potential of the past marine microbiome (approximately 300 years old) inhabiting under the ice shelf.

**Figure 4.**
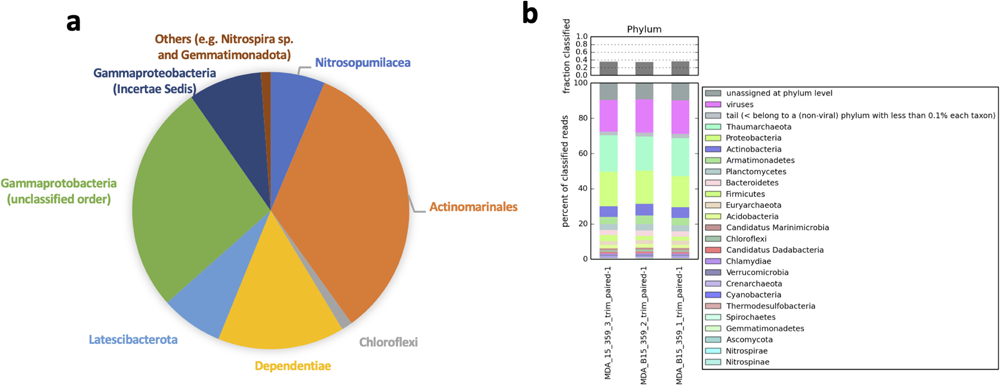
Metagenomics and 16S rRNA gene information of microbial DNA preserved in the marine ice core B15. (a) Taxonomic assignment of 16S rRNA reads extracted with *miTaq* protocol from metagenomic data. Remarkably, as with the 16S rRNA gene sequences recovered from PCR of multiple-displacement amplification (MDA) products, no common contaminant sequences were found from molecular reagents, human skin or other exogenous sources. (b) Taxonomic assignment of all raw reads obtained in metagenomes (triplicate samples) from the marine ice core B15 dating 275-328 years old before drilling.

**Figure 5.**
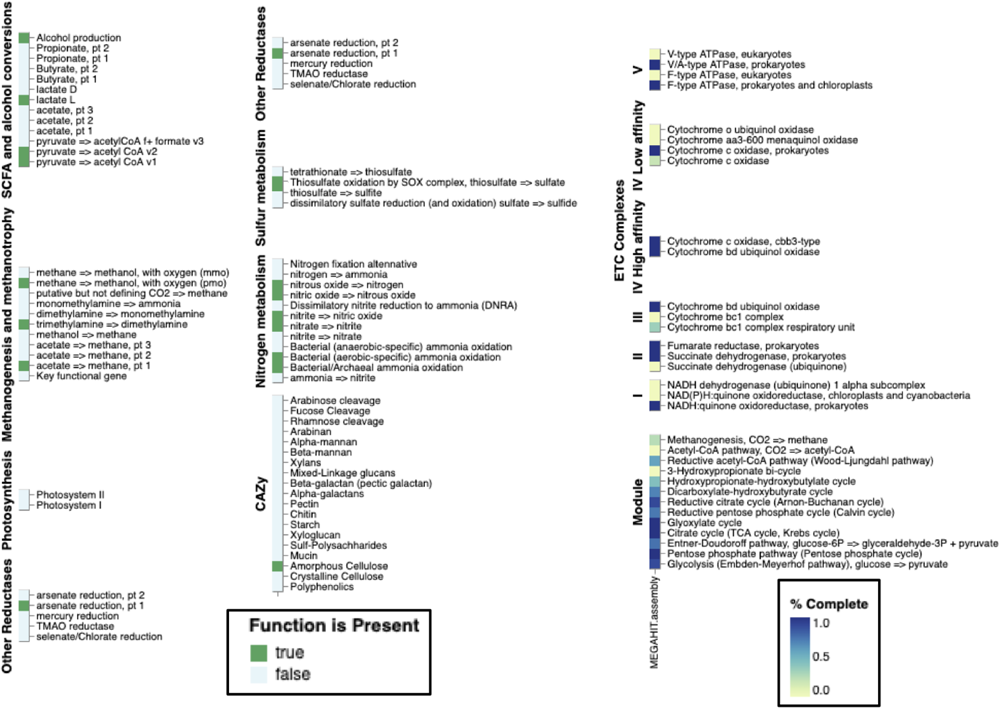
Metabolic potential reconstructed from microbial DNA preserved in the analyzed marine ice. Metagenome samples were assembled with Megahit and Spades (single-cell option) assemblers. Gene prediction and annotation was carried out with DRAM annotator. Functional categories involved in key metabolic pathways are shown.

Although most of the annotated genes indicate that aerobic metabolism predominates (Fig. 5), a few metabolic marker genes involved in methanogenesis and sulphate respiration including 16S rRNA gene sequences related to microbes found in deep marine sediments and cold permafrost (Fig. S8) were also detected. Likely these sequences come from particle-attached microbes originally transported from sediments of the grounding line of the ice shelf that were later frozen during marine ice formation. It is important to remind that beyond the Filchnner Ronne Ice Shelf, as in all Antarctic ice shelves, there is a strong cavity circulation current tightly inter-connected with large-scale atmospheric and oceanic circulation patterns^55^. Metagenome-assembled genomes (MAGs) from these samples representing key microbes were also retrieved complementing our genetic dataset (Table S2). A total of 10 medium-quality MAGs and 1 high-quality MAG (quality threshold according to^56^) were obtained belonging to different phyla (Desulfobacterota, Dependentiae, Actinobacteriota, Proteobacteria, and Chlamydiota; Supplementary Table 2). Genomic analysis with the nearest genome in GTDB-k showed that most of the retrieved MAGs were distantly related (data not shown).

Remarkably, despite the limited dataset generated in this pilot proof-of-concept study, we were able to calibrate the evolutionary genomic and genetic changes of *Nitrosopumilus spp.* preserved in the marine ice core 300 years ago, by comparing our data with those *Nitrosopumilus spp.* recently collected from seawater under an Antarctic ice shelf^30^. To achieve this, we estimated the rate of genomic SNPs over time and the evolutionary divergence of *amoA* gene (i.e. substitution rate over time) (Fig. 6). Data for *amoA* gene suggest that approximately 100 years are required for ≍1 amino acid (aa) substitution per each 100 aa positions of the protein (Fig. 6B). The obtained rate of genomic SNPs for *Nitrosopumilus* spp. was about 2,500 accumulated SNPs per 1 Mb and 100 years (Fig. 6A). Whether these evolutionary changes remained constant over the last 300 years or accelerated during post-industrial periods remains an open question. This will be properly addressed when a more complete dataset of samples and periods is analyzed.

**Figure 6.**
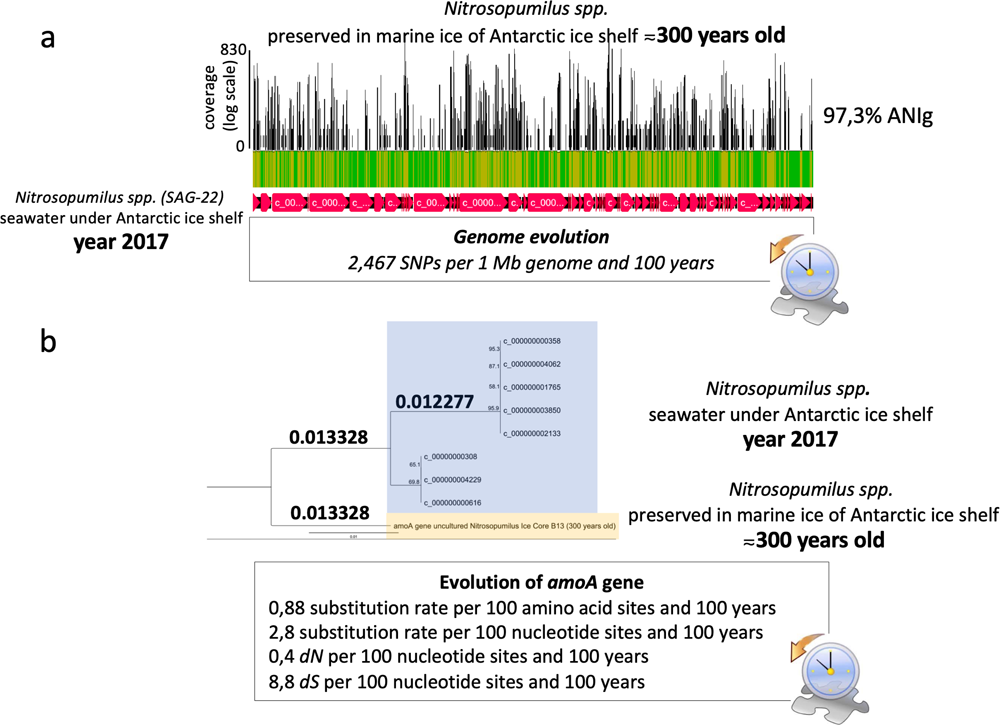
Calibration of evolutionary changes over time for a representative microbe preserved in the marine ice core B15. (a) Genome mapping of metagenomic reads obtained from the marine ice core B15 against *Nitrosopumilus* spp. genome (SAG-22) collected from seawater in 2017 right beneath an Antarctic ice shelf. This comparison between samples taken more than 300 years apart allow us to calibrate rate of genomic SNPs (A) and the substitution rate of selected genes, such as *amoA* (B). Similar phylogenetic tree was obtained with different evolutionary models. Read mapping was performed with Bowttie and SNPs calculated

Finally, although we cannot totally rule out some potential contamination from ice core surface through ice cracking or manipulation, the metagenomic data obtained here after a careful decontamination indicate that microbes are indeed of a typical marine source (i.e. marine Thaumarchaea or Actinomarinales; see Fig. 4 and Fig. S8 and S10). It is important to remark that we have not found sequences of common abundant bacteria present in the Antarctic surface snow, which are typically *Flavobacterium, Hydrogenophaga, Ralstonia, Janthinobacterium, Caulobacter*, and *Pseudomonas*^57^. Furthermore, contamination signal from resistant airborne microbes over Antarctica (e.g. *Bacillus* spp. and fungi)^58^ or from skin microbes that might be introduced during drilling and later manipulation were not observed. Finally, it is important to clarify that during the drilling campaign in 1992 to collect this marine ice core studied here, the drilling did not reach the bottom of the ice shelf but it stopped several meters away from the basal part of the ice shelf, preventing thus the entry and potential contamination from marine microbes naturally present in the seawater under the Filchnner-Ronne Ice Shelf. Thus, although dealing with ice cores is technically complex and always challenging to show and demonstrate the data validity, here, data suggest that genetic information obtained from the inner part of the ice core (1 ml) seem to actually represent preserved microbial DNA from the past ocean.

In conclusion, our study demonstrates that in the marine ice beneath the Antarctic ice shelves, there is a potential genuine microbial DNA stored in different layers of antiquity that can be dated using robust and reliable ice-flow models. Remarkably, our data suggest that this novel approach successfully recovered past marine, microbial DNA trapped in marine ice under the Antarctica and open the possibility to “travel to the past”. It is important to underscore that although our study does not allow to address a comprehensive and fine evolutionary analysis within the last period of 300 years because of the limited number of available samples, our approach was indeed able to calibrate genetic and genomic evolutionary changes for a selected key microbe that can extended for other groups and functions. Our proof-of-concept study therefore initiates a novel avenue within the field of climate change research, allowing for an exploration of how anthropocentric pressure has potentially shaped microbial communities over time. Although the data retrieved here from microbes frozen >300 years ago, prior to the dramatic CO_2_ increase, represent a valuable baseline for further genetic comparison, the results should be complemented with a more complete dataset of marine ice samples from different ice cores and different time periods to obtain reliable outcomes. Furthermore, radioisotopic assays for ice-core dating could be taken into considerations when a fine-scale time resolution of marine ice is desired. To expand these genomic databases, additional Antarctic ice shelf drilling campaigns are warranted. Currently, the availability of marine ice samples from Antarctic ice shelves is very limited, and this scarcity is expected to worsen in the future due to the accelerating rate of ice melting caused by climate change.

## Materials and methods

### Age–depth estimation of ice core

Ice core B15 used in this study was retrieved from a hole (65-72 mm diameter) drilled mechanically in 1992 on the Ronne Ice Shelf (Antarctic; 76° 58’ S 52° 16′ W) with the total length of 320.7 m during the cruise ANT VIII/5 of RV Polarstern^29^. The upper 152.8 m consists of meteoric ice followed by a 167.9 m thick column of marine ice that has been aggregated during the time the glacier flow takes from the grounding line to the location of B15 at the front of the Ronne Ice Shelf. In order to estimate the age of the aggregated marine ice at B15, we analyzed basal melt and accretion rates by Adusumilli et al.^42^ (data from http://library.ucsd.edu/dc/object/bb0448974g) along the flowline, accounting for the dynamic thinning from vertical strain (see ref ^43^ and Fig. S2-S4). Based on the MEaSUREs ice flow velocities ^59,60^, we computed both the flowline and the flow duration from the grounding line. For each 250 m long segment of the flowline, we calculated the change in marine ice thickness (Δ*H*_*m*_)

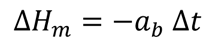

from the accretion or melting (*a*_*b*_, positive for melting) during the flow duration (Δ*t*). By accounting for the dynamic thinning from the vertical strain rate (*Ė_zz_*), we obtain the marine ice thickness (*H*_*m*_) as a function of the time step (*i*):

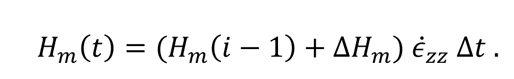

The resulting marine ice thickness evolution along the flowline of B15 is shown in Fig. S3. Based on this estimation we obtained a marine ice thickness of 177.5 m, which is close to the observed marine ice thickness of 167.9 m in the retrieved ice core.

In order to obtain the age–depth estimation, we stored the thickness of the accumulated marine ice Δ*H*_*m*_ as a new *n* = *n* + 1-th layer to an array ℎ_*m*_,

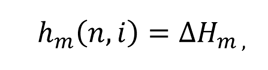

and the age *τ* of the layer

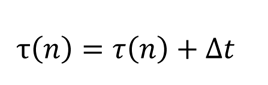

in case of accretion. If no accretion (or melting) occurred (Δ*H*_*m*_ ≤ 0), we reduced the thickness of the *n*-th layer:

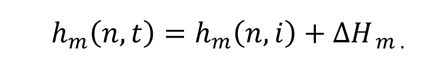

In case the amount of melted marine ice exceeded the thickness of the *n*-th layer (ℎ_*m*_(n, i) ≤ Δ*H*_*m*_), we removed this ℎ_*m*_(*n*, *i*) = [] as well as the age *τ*(*n*) = [] and also reduced the ice thickness of the *n* − 1-th layer by the remaining difference.

After each segment, we considered the dynamic thinning of every layer (1: *n*):

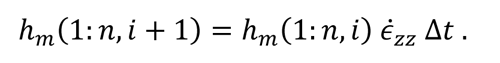

According to this analysis, the oldest marine ice at B15 is 1100 years old and the marine ice at 134,7 m below the meteoric ice used in this study, was accumulated 275 to 328 years ago (Fig. S4). The age uncertainty takes the reported uncertainty from satellite remote sensing methods for estimates of basal melt/accretion rates into account. By applying this method, we assume that the ice flow velocities, strain rates and especially basal melt/accretion rates, which we observed in the last couple of years, are representative for the last thousand years and more. However, the evolution of the quantities over this time period is not quantified and so we cannot express this uncertainty in our analysis.

### Ice core processing and decontamination method

Ice core B15 was properly stored at the Alfred Wegener Institute (AWI) located in Bremerhaven (Germany). In 2022, ice core B15 was cut in pieces of 80 cm length approximately and in June 2022, samples were shipped to the University of Alicante under freezing conditions. Upon arrival, samples were conserved at −24°C until use.

In our study, we applied the previously described method for decontaminating ice core samples used for microbiology^9,61^ with very minor modifications. Samples processed from the ice core B15 were from layers corresponding to approximately 277-329 years old before drilling (January 1992). Different independent inner ice core samples (see Supplementary Table 1) were extracted and processed for metagenomics to check for variability between the same 80 cm ice core piece. All sample processing (Fig. S1) was carried out within an area with a dedicated laminar flow cabin used for single-cell genomic applications. The cabin was thoroughly bleached and UV for 30 min as described^41,62^ before starting the process. All the material used for manipulating the ice core, such as that for cutting or scrapping was autoclaved and UV for 15 min before use. Ice core samples were precisely cut at the required size using a 100-240V /18W Styrofoam Cutting tool with 18 cm bow hot blade and Electronic Voltage Transformer Adaptor, brand Gochange in sterile condition in a special dedicated cabin used only for single-cell and virus genomic applications. Then, surface of the ice core sample was scratched (≍5 mm), and immediately the ice sample was covered and washed with 70% freshly prepared ethanol with sterilized mQ water followed by another washing step with mQ water as indicate by Zhong and colleagues^9,61^. Finally, mQ water was let run-off from the inner ice core sample for a minute, and sample was thawed at 10°C in a dedicated sterile container within the laminar flow cabin. During the whole processing, manipulators and researchers had individual protection equipment (caps, glasses, gloves, mask, etc…). Gloves were frequently bleached as in single cell genomic protocols^41,62^.

### Fluorescence Microscopy

Samples for microscopy were obtained from the inner ice core previously decontaminated and thawed (Fig. S1). Sample was fixed in 2% w/w formaldehyde (Sigma) at 4°C for 2 h. For confocal and epifluorescence microscopy, melted inner ice core sample were stained with DAPI 1μg/ml final concentration for 5 min, SYBR Gold 25x from commercial stock for 15 min, and Live/Dead BacLight^™^ kit (Invitrogen) containing propidium iodide and SYTO9 according to manufactureŕs protocol. These dyes are DNA-specific and very common in Microbiology^62–65^. For DAPI staining, a total of 10 ml of fixed sample was filtered in 0,22 μm pore filters Isopore^TM^ Policarbonate Membrane Filters, while for SYBR Gold staining, a total of 4 ml of sample was filtered in 0.02 μm Watman® Anodisc filter. For the Live/Dead BacLight^™^ staining kit propidium, a total of ≍200 ml of fixed sample was concentrated to 1 ml with centrifugal filtering Systems Amicon^®^ 10 KDa. Then, 100 ul of sample was stained as manufacturer’s protocol and filtered in a small region of a 0,02 μm pore size Whatman® Anodisc filter, delimited with a PAP-Pen (2 mm tip). In addition, unstained fixed sample (4ml) was filtered in 0.02 mm Watman® Anodisc filter to check for potential autofluorescence. For all samples prepared for microscopy, antifading reagent Citifluor AF1 (Electron Microscopy Science company) was added on filters (≍10-20 μl) and covered with a cover slip. Samples were inspected under a Zeiss LSM 800 Confocal Laser Scanning Microscope and in an epifluorescence microscope Leica DM4000 B equipped with a camera Leica Flexacam C3. For confocal microscopy analysis, lasers were set to 10% intensity and photomultiplier power detectors to 650V. Unstained samples were inspected in all color spectrum and excited with UV, blue, green and red lasers and processed as samples.

### Electron scanning microscopy

Thawed inner ice core sample (volume 10 ml) was fixed with 2% glutaraldehyde during 1 h and filtered with 0,05 μm SPI-Pore^TM^ Polycarbonate Track Etch Membrane Filters as described in^66^ for SEM. A total of 1,5 or 5 ml sample volume was filtered. Samples were visualized in a high-resolution scanning electron microscope Jeol model IT500HR/LA with EDS analysis.

### DNA extraction and optimization of single cell technologies protocol for Antarctic Ice Cores

Different ice core samples were processed from the same marine ice piece of 275-328 years old before drilling (named as PltCore_1 and PltCore_2; see Fig. S1) to obtain independent samples and sequencing replicates. Melted samples (different volumes from 300 to 1000 ml) were filtered on 0,22 μm pore size hydrophilic PVDF filters Durapore^®^ and DNeasy Power Soil Pro^®^ Kit (Qiagen) was applied for DNA extractions as is the most common and efficient method used in microbiome surveys^51,67,68^. Eluate volume samples from 0,22 μm pore size hydrophilic PVDF filters Durapore^®^ containing potentially biological nanoparticles, such as viruses, were also collected and filtered on 0,02μm pore size Anodisc inorganic filters Whatman^®^. DNA extraction from the fraction filtered through 0,02 μm pore size Anodisc inorganic filters was carried out with QIAamp MinElute Virus Kit^®^. In any case, Qubit^TM^ fluorimeter using the high sensitivity kit (HS) could detect DNA despite repeating the experiment several times and scaling up the volume up to more than 1 L. Same results were obtained when other DNA extraction kits were tested, such as Circulating DNA Purification Mini Spin^®^ Kit (Canvax), BS Buccal Saliva Genomic DNA Extraction^®^ Kit (Canvax), Blood Genomic DNA Extraction^®^ Kit (Canvax) and DNeasy Power Waste^®^ Kit (Qiagen). In addition, a classic protocol of DNA extraction based on phenol:chloroform:isoamyl alcohol and alcohol precipitation was unsuccessfully tried for ≍200 ml of melted sample. A direct precipitation of free DNA present in ≍200 ml of melted sample was also tried without any success with isoamyl alcohol procedure. All these common methods failed to retrieve measurable DNA since fluorometric measurements of the available microbial DNA (e.g. high molecular weight) with Qubit high sensitivity kit was below the limit. Bear in mind that for the whole ice core B15 (65-72 mm diameter; a few dozen meter length spanning different periods), there is only a few liters of available marine ice for the research community. Since the amount of volume of sample is very limited and precious and data suggested very tiny amount of available DNA in samples for molecular analysis, we opted for implementing single cell genomic technologies to extract and obtain the genetic information from inner ice core samples.

The single cells genomics methods is able to extract, amplify and sequence the genome from as little as one single viral particle or cell^62,69–73^. Here, we implemented the single-cell and -virus genomic protocols with very little modification. As a starting material, first a few milliliters (<5 ml) from the inner ice core previously decontaminated and thawed were used for single cell genomics. A total of 0,6 μl of melted ice core was pipetted in each one of the well in a 384-well plate. To ensure that we capture and represented properly in our experiments the microbial diversity given the limited amount of marine ice sample, other pieces with different sizes and volumes (up to 200 ml approximately) from the inner ice core were concentrated with Amicon® 10 KDa and then the ultraconcentrated volume used as input for single-cell genomics as well (0,6 μl of sample per well). Then, subsequent cell lysis and real time multiple-displacement amplification (MDA) was performed as in many previous single cell and virus genomic surveys^39,62,70,74,75^. In brief, to 0.6 ul of thawed sample in each well, we added 0.7 uL of lysis buffer D2 (Qiagen) and after 5 min, pH was neutralized employing 0.7 uL of Stop solution (Qiagen). Then, whole-genome amplification was carried out at 45°C by real time MDA using the novel Equiphi 29 enzyme as described^74^ (New England Biolab) for approximately 3-4 h including blanks (no sample added) and positive controls (0.6 ng of internal DNA as in^70^) to monitor DNA contamination in reagents or DNA introduced by manipulation. Real-time MDA was monitored by fluorescence thanks to SYTO-9 dye that was added to the MDA reaction, allowing the identification of positive amplification during MDA. Common procedures used during single cell genomics previously, such as decontamination of DNA polymerase and reagents, described were also used in our experiments (see more details in^41,62^).

### Sequencing and data analysis

Triplicate libraries and sequencing from the same sample (PlteCore_1) was performed, while a single library and sequencing reaction was performed for *PltCore_2*. For all samples, paired-end Illumina® DNA Prep, (M) Tagmentation with IDT® for Illumina® DNA/RNA UD Indexes was performed according to manufacturer’s protocol and sequenced in a Hiseq sequencer (150x2 PE) by Macrogen company (Korea).

The read sequences were trimmed with Trimmomatic v.0.36^76^, changing the default parameter of sliding windows minimum quality to 30 and minimum read length to 50, and assessed with FastQC v.0.12.1 according to default parameters (https://www.bioinformatics.babraham.ac.uk/publications.html). The diversity of every plate and bulk was assessed with Kaiju v.1.7.3^77^., using NCBI BLASTnr+euk as the reference database and changing the default abundance filter to 0,1. Nonpareil tool from Kostas Lab^54,78^ was used to check the sample diversity and compared each other with Mash metagenome distance^79^ (Suppl. Table 1), using forward trimmed sequences as input. Program mTAG^53^ was applied to the forward trimmed reads for each sample to recover 16S rRNA gene classification. In parallel, 16S rRNA gene PCR and Sanger sequencing was performed from 60 randomly amplified wells obtained in MDA using 27F Bacteria16S rRNA primer (5’-AGA GTT TGA TCM TGG CTC AG-3’) and 907R Bacteria16S rRNA primer (5’-CCG TCA ATT CMT TTG AGT TT-3’). Primers were obtained from IDT company. The obtained sequences were quality trimmed with Geneious^80^. After that, Blast analysis was done against nr database of NCBI Genbank and compared with SILVA^81,82^ and RDP^83^ database.

Trimmed reads from Illumina Hight Genomic Sequencing were assembled with Spades^75,84^ v 3.15.3 (using the Single Cell option) and quality assessed with Quast^85^. Contigs were annotated with Dram program^86,87^ using minimum contig length 2500 bp and bit score threshold of 60.

From the assembly, also different binning programs were used as Metabat2 v1.7^88^, Concoct v1.1^89^ and MaxBin2 v2.2.4^90^, using in all cases a minimum contig length of 2500 bp. All the bins obtained were assessed with Ckeck M^91^ and optimized with DAS tools binning^92^, using a score threshold 0,5, duplicate penality 0,6 and megabin penality 0,5. After that, the selected bins were classified with GTDB-Tk database^93^ and the nearest genome in GTDB-Tk was selected for average nucleotide identity comparison with Jspecies program^94^. Those bins showing more than 10% of contamination were discarded. Several bioinformatics analysis were performed online with Kbase^95^. Read mapping was performed with Bowtie2 program (default parameters)^96^ implemented in Geneious bioinformatic package version R9.0^80^. For that, quality trimmed reads obtained from sample Plt_Core_1 were mapped against the genome of *Nitrosopumilus* spp. (SAG-22) obtained in 2017 from seawater under an Antarctic ice shelf^30^ and also against the isolated strain *Nitrosopumilus maritimus SCM1. amoA* genes belonging to different *Nitrosopumilus* SAGs (SAGs no. 22, 24, 28, 4, 40, 57, 61, 64, 8) and one MAG (MAG-12) obtained in 2017 from seawater under an Antarctic ice shelf^30^ were aligned along with *amoA* sequence of *Nitrosopumilus* spp. obtained in the ice core B15 dated 300 years old. Alignment was performed with ClustalW implemented in Geneious bioinformatic package version R9.0^80^. Phylogenetic tree was calculated with Jukes-Cantor distance model (neighbour-joining tree build method) with a bootstrapping of 1,000. Number of SNPs were calculated as described^40^.

## Acknowledgements

We thank the research grants funded by the Spanish Ministry of Science and Innovation and Agencia Estatal de Investigación (PID2021-125175OB-I00).

## Author contribution

## Additional information

**Supplementary information.** The online version contains supplementary material

## Data accession

Illumina sequencing data is deposited under SRA accession number PRJNA978593

## References

1. Breitbart, M., Bonnain, C., Malki, K. & Sawaya, N. A. Phage puppet masters of the marine microbial realm. Nat. Microbiol. 1 (2018) doi:10.1038/s41564-018-0166-y.

2. Roux, S. et al. Ecogenomics and potential biogeochemical impacts of globally abundant ocean viruses. Nature 537, 689–693 (2016).

3. Del Campo, J., Not, F., Forn, I., Sieracki, M. E. & Massana, R. Taming the smallest predators of the oceans. ISME J. 7, 351–358 (2012).

4. Not, F., del Campo, J., Balagué, V., de Vargas, C. & Massana, R. New insights into the diversity of marine picoeukaryotes. PLoS One 4, e7143 (2009).

5. Hutchins, D. A. & Fu, F. Microorganisms and ocean global change. Nat. Microbiol. 2017 26 2, 1–11 (2017).

6. Hutchins, D. A. et al. Climate change microbiology — problems and perspectives. Nat. Rev. Microbiol. 17, 391–396 (2019).

7. Vaqué, D. et al. Warming and CO2 Enhance Arctic Heterotrophic Microbial Activity. Front. Microbiol. 10, 494 (2019).

8. Cavicchioli, R. et al. Scientists’ warning to humanity: microorganisms and climate change. Nat. Rev. Microbiol. 17, (2019).

9. Zhong, Z.-P. et al. Glacier ice archives nearly 15,000-year-old microbes and phages. Microbiome 2021 91 **9**, 1–23 (2021).

10. Zhang, R., Weinbauer, M. G. & Peduzzi, P. Aquatic Viruses and Climate Change. Curr. Issues Mol. Biol. 41, 357–380 (2021).

11. Evans, C. & Brussaard, C. P. D. Regional Variation in Lytic and Lysogenic Viral Infection in the Southern Ocean and Its Contribution to Biogeochemical Cycling. Appl. Environ. Microbiol. 78, 6741–6748 (2012).

12. Biggs, T. E. G., Huisman, J. & Brussaard, C. P. D. Viral lysis modifies seasonal phytoplankton dynamics and carbon flow in the Southern Ocean. ISME J. 1–8 (2021) doi:10.1038/s41396-021-01033-6.

13. Lara, E. et al. Experimental evaluation of the warming effect on viral, bacterial and protistan communities in two contrasting Arctic systems. Aquat. Microb. Ecol. 70, 17–32 (2013).

14. Newbold, L. K. et al. The response of marine picoplankton to ocean acidification. Environ. Microbiol. 14, 2293–307 (2012).

15. Malits, A. et al. Viral-Mediated Microbe Mortality Modulated by Ocean Acidification and Eutrophication: Consequences for the Carbon Fluxes Through the Microbial Food Web. Front. Microbiol. 12, (2021).

16. Das, S. & Mangwani, N. Ocean acidification and marine microorganisms: responses and consequences. Oceanologia 57, 349–361 (2015).

17. Boyd, P. W. et al. Experimental strategies to assess the biological ramifications of multiple drivers of global ocean change-A review. Glob. Chang. Biol. 24, 2239–2261 (2018).

18. Vaqué, D. et al. Warming and CO2 enhance arctic heterotrophic microbial activity. Front. Microbiol. 10, 1–13 (2019).

19. Vaqué, D. et al. Warming and CO2 Enhance Arctic Heterotrophic Microbial Activity. Front. Microbiol. 10, 494 (2019).

20. Thomas, M. K., Kremer, C. T., Klausmeier, C. A. & Litchman, E. A global pattern of thermal adaptation in marine phytoplankton. Science *(80-.).* **338**, 1085–1088 (2012).

21. Bopp, L. et al. Multiple stressors of ocean ecosystems in the 21st century: Projections with CMIP5 models. Biogeosciences 10, 6225–6245 (2013).

22. Ibarbalz, F. M. et al. Global Trends in Marine Plankton Diversity across Kingdoms of Life. Cell 179, 1084–1097.e21 (2019).

23. Beaugrand, G. et al. Prediction of unprecedented biological shifts in the global ocean. Nat. Clim. Chang. 2019 93 **9**, 237–243 (2019).

24. Frémont, P. et al. Restructuring of plankton genomic biogeography in the surface ocean under climate change. Nat. Clim. Chang. 2022 1–9 (2022) doi:10.1038/s41558-022-01314-8.

25. Pinsky, M. L., Worm, B., Fogarty, M. J., Sarmiento, J. L. & Levin, S. A. Marine taxa track local climate velocities. Science *(80-.).* **341**, 1239–1242 (2013).

26. Barton, A. D., Irwin, A. J., Finkel, Z. V. & Stock, C. A. Anthropogenic climate change drives shift and shuffle in North Atlantic phytoplankton communities. Proc. Natl. Acad. Sci. U. S. A. 113, 2964–2969 (2016).

27. Adusumilli, S., Fricker, H. A., Medley, B., Padman, L. & Siegfried, M. R. Interannual variations in meltwater input to the Southern Ocean from Antarctic ice shelves. Nat. Geosci. 2020 139 **13**, 616–620 (2020).

28. Rignot, E., Jacobs, S., Mouginot, J. & Scheuchl, B. Ice-shelf melting around antarctica. Science *(80-.).* **341**, 266–270 (2013).

29. Oerter, H. et al. Evidence for basal marine ice in the Filchner–Ronne ice shelf. Nat. 1992 3586385 **358**, 399–401 (1992).

30. Martínez-Pérez, C. et al. Phylogenetically and functionally diverse microorganisms reside under the Ross Ice Shelf. Nat. Commun. 2022 131 **13**, 1–15 (2022).

31. Craven, M. et al. Borehole imagery of meteoric and marine ice layers in the Amery Ice Shelf, East Antarctica. J. Glaciol. 51, 75–84 (2005).

32. Joughin, I. & Vaughan, D. G. Marine ice beneath the Filchner-Ronne Ice Shelf, Antarctica: A comparison of estimated thickness distributions. Ann. Glaciol. 39, 511–517 (2004).

33. Tison, J. L., Khazendar, A. & Roulin, E. A two-phase approach to the simulation of the combined isotope/salinity signal of marine ice. J. Geophys. Res. Ocean. 106, 31387– 31401 (2001).

34. Thomas, D. N. & Dieckmann, G. S. Antarctic Sea Ice--a Habitat for Extremophiles. Science *(80-.).* **295**, 641–644 (2002).

35. Antony, R. et al. Diversity and physiology of culturable bacteria associated with a coastal Antarctic ice core. Microbiol. Res. 167, 372–380 (2012).

36. Adusumilli, S., Fricker, H. A., Medley, B., Padman, L. & Siegfried, M. R. Interannual variations in meltwater input to the Southern Ocean from Antarctic ice shelves. Nat. Geosci 2020 139 **13**, 616–620 (2020).

37. Griggs, J. A. & Bamber, J. L. Antarctic ice-shelf thickness from satellite radar altimetry. J. Glaciol. 57, 485–498 (2011).

38. Eicken, H., Oerter, H., Miller, H., Graf, W. & Kipfstuhl, J. Textural characteristics and impurity content of meteoric and marine ice in the Ronne Ice Shelf, Antarctica. J. Glaciol. 40, 386–398 (1994).

39. Martínez Martínez, J., Martinez-Hernandez, F. & Martinez-Garcia, M. Single-virus genomics and beyond. Nat. Rev. Microbiol. 18, 705–716 (2020).

40. Martinez-Hernandez, F. et al. Single-virus genomics reveals hidden cosmopolitan and abundant viruses. Nat. Commun. 8, 15892 (2017).

41. Woyke, T. et al. Decontamination of MDA reagents for single cell whole genome amplification. PLoS One 6, e26161 (2011).

42. Adusumilli, S., Fricker, H. A., Medley, B., Padman, L. & Siegfried, M. R. Interannual variations in meltwater input to the Southern Ocean from Antarctic ice shelves. Nat. Geosci. 13, 616–620 (2020).

43. Alley, K. E. et al. Continent-wide estimates of Antarctic strain rates from Landsat 8-derived velocity grids. J. Glaciol. 64, 321–332 (2018).

44. Van Liefferinge, B. & Pattyn, F. Using ice-flow models to evaluate potential sites of million year-old ice in Antarctica. Clim. Past 9, 2335–2345 (2013).

45. Humbert, A. et al. On the evolution of an ice shelf melt channel at the base of Filchner Ice Shelf, from observations and viscoelastic modeling. Cryosphere 16, 4107–4139 (2022).

46. Licciulli, C. et al. A full Stokes ice-flow model to assist the interpretation of millennial-scale ice cores at the high-Alpine drilling site Colle Gnifetti, Swiss/Italian Alps. J. Glaciol. 66, 35–48 (2020).

47. Grinsted, A., Moore, J., Spikes, V. B. & Sinisalo, A. Dating Antarctic blue ice areas using a novel ice flow model. Geophys. Res. Lett. 30, (2003).

48. Belilla, J. et al. Active Microbial Airborne Dispersal and Biomorphs as Confounding Factors for Life Detection in the Cell-Degrading Brines of the Polyextreme Dallol Geothermal Field. MBio 13, (2022).

49. Klauth, P., Wilhelm, R., Klumpp, E., Poschen, L. & Groeneweg, J. Enumeration of soil bacteria with the green fluorescent nucleic acid dye Sytox green in the presence of soil particles. J. Microbiol. Methods 59, 189–198 (2004).

50. Minich, J. J. et al. KatharoSeq Enables High-Throughput Microbiome Analysis from Low-Biomass Samples. mSystems 3, e00218–17 (2018).

51. Knight, R. et al. Best practices for analysing microbiomes. Nat. Rev. Microbiol. 16, 410– 422 (2018).

52. Gawad, C., Koh, W. & Quake, S. R. Single-cell genome sequencing: current state of the science. Nat. Rev. Genet. 17, 175–188 (2016).

53. Sunagawa, S. et al. Structure and function of the global ocean microbiome. Science *(80-.).* **348**, 1–9 (2015).

54. Rodriguez-r, L. M. & Konstantinidis, K. T. Nonpareil: a redundancy-based approach to assess the level of coverage in metagenomic datasets. Bioinformatics 30, 629–35 (2014).

55. Hattermann, T. et al. Observed interannual changes beneath Filchner-Ronne Ice Shelf linked to large-scale atmospheric circulation. Nat. Commun. 2021 121 **12**, 1–11 (2021).

56. Bowers, R. M. et al. Minimum information about a single amplified genome (MISAG) and a metagenome-assembled genome (MIMAG) of bacteria and archaea. Nat. Biotechnol. 35, 725–731 (2017).

57. Lopatina, A. et al. Metagenomic analysis of bacterial communities of antarctic surface snow. Front. Microbiol. 7, 179402 (2016).

58. Pearce, D. A. et al. Microorganisms in the atmosphere over Antarctica. FEMS Microbiol. Ecol. 69, 143–157 (2009).

59. Mouginot, J., Rignot, E. & Scheuchl, B. Continent-Wide, Interferometric SAR Phase, Mapping of Antarctic Ice Velocity. Geophys. Res. Lett. 46, 9710–9718 (2019).

60. MEaSUREs Phase-Based Antarctica Ice Velocity Map, Version 1 | National Snow and Ice Data Center. https://nsidc.org/data/nsidc-0754/versions/1.

61. Zhong, Z.-P. et al. Clean Low-Biomass Procedures and Their Application to Ancient Ice Core Microorganisms. Front. Microbiol. 9, (2018).

62. Rinke, C. et al. Obtaining genomes from uncultivated environmental microorganisms using FACS-based single-cell genomics. Nat. Protoc. 9, 1038–48 (2014).

63. Porter, K. G. & Feig, Y. S. The use of DAPI for identifying and counting aquatic microflora1. Limnol. Oceanogr. 25, 943–948 (1980).

64. Chen, F., Lu, J. R., Binder, B. J., Liu, Y. C. & Hodson, R. E. Application of digital image analysis and flow cytometry to enumerate marine viruses stained with SYBR gold. Appl. Environ. Microbiol. 67, 539–45 (2001).

65. Nocker, A., Richter-Heitmann, T., Montijn, R., Schuren, F. & Kort, R. Discrimination between live and dead cells in bacterial communities from environmental water samples analyzed by 454 pyrosequencing. Int. Microbiol. 13, 59–65 (2010).

66. Golding, C. G., Lamboo, L. L., Beniac, D. R. & Booth, T. F. The scanning electron microscope in microbiology and diagnosis of infectious disease. Sci. Reports 2016 61 **6**, 1–8 (2016).

67. Eisenhofer, R. et al. Contamination in Low Microbial Biomass Microbiome Studies: Issues and Recommendations. Trends Microbiol. 27, 105–117 (2019).

68. Bolyen, E. et al. Reproducible, interactive, scalable and extensible microbiome data science using QIIME 2. Nat. Biotechnol. 37, 852–857 (2019).

69. Allen, L. Z. et al. Single virus genomics: a new tool for virus discovery. PLoS One 6, e17722 (2011).

70. Martinez-Hernandez, F. et al. Single-virus genomics reveals hidden cosmopolitan and abundant viruses. Nat. Commun. 8, 15892 (2017).

71. Martínez Martínez, J., Martinez-Hernandez, F. & Martinez-Garcia, M. Single-virus genomics and beyond. Nat. Rev. Microbiol. 2020 1812 18, 705–716 (2020).

72. Wilson, W. H. et al. Genomic exploration of individual giant ocean viruses. ISME J. 11, 1736–1745 (2017).

73. Ghylin, T. W. et al. Comparative single-cell genomics reveals potential ecological niches for the freshwater acI Actinobacteria lineage. ISME J. 8, (2014).

74. Stepanauskas, R. et al. Improved genome recovery and integrated cell-size analyses of individual uncultured microbial cells and viral particles. Nat. Commun. 8, 1–10 (2017).

75. Swan, B. K. et al. Potential for chemolithoautotrophy among ubiquitous bacteria lineages in the dark ocean. Science 333, 1296–300 (2011).

76. Bolger, A. M., Lohse, M. & Usadel, B. Trimmomatic: a flexible trimmer for Illumina sequence data. Bioinformatics 30, 2114–2120 (2014).

77. Menzel, P., Ng, K. L. & Krogh, A. Fast and sensitive taxonomic classification for metagenomics with Kaiju. Nat. Commun. 7, 11257 (2016).

78. Rodriguez-r, L. M. & Konstantinidis, K. T. Sequence analysis Nonpareil : a redundancy-based approach to assess the level of coverage in metagenomic datasets. 30, 629–635 (2014).

79. Ondov, B. D. et al. Mash: fast genome and metagenome distance estimation using MinHash. Genome Biol. 17, 132 (2016).

80. Kearse, M. et al. Geneious Basic: an integrated and extendable desktop software platform for the organization and analysis of sequence data. Bioinformatics 28, 1647–9 (2012).

81. Klindworth, A. et al. Evaluation of general 16S ribosomal RNA gene PCR primers for classical and next-generation sequencing-based diversity studies. Nucleic Acids Res. 41, e1 (2013).

82. Quast, C. et al. The SILVA ribosomal RNA gene database project: improved data processing and web-based tools. Nucleic Acids Res. 41, D590–6 (2013).

83. Wang, Q., Garrity, G. M., Tiedje, J. M. & Cole, J. R. Naive Bayesian classifier for rapid assignment of rRNA sequences into the new bacterial taxonomy. Appl. Environ. Microbiol. 73, 5261–7 (2007).

84. Bankevich, A. et al. SPAdes: a new genome assembly algorithm and its applications to single-cell sequencing. J. Comput. Biol. 19, 455–77 (2012).

85. Gurevich, A., Saveliev, V., Vyahhi, N. & Tesler, G. QUAST: quality assessment tool for genome assemblies. Bioinformatics 29, 1072–5 (2013).

86. Singleton, C. M. et al. Connecting structure to function with the recovery of over 1000 high-quality metagenome-assembled genomes from activated sludge using long-read sequencing. Nat. Commun. 12, 2009 (2021).

87. Shaffer, M. et al. DRAM for distilling microbial metabolism to automate the curation of microbiome function. Nucleic Acids Res. 48, 8883–8900 (2020).

88. Lin, H.-H. & Liao, Y.-C. Accurate binning of metagenomic contigs via automated clustering sequences using information of genomic signatures and marker genes. Sci. Rep. 6, 24175 (2016).

89. Alneberg, J. et al. Binning metagenomic contigs by coverage and composition. Nat. Methods 11, 1144–1146 (2014).

90. Wu, Y.-W., Simmons, B. A. & Singer, S. W. MaxBin 2.0: an automated binning algorithm to recover genomes from multiple metagenomic datasets. Bioinformatics 32, 605–607 (2016).

91. Parks, D. H., Imelfort, M., Skennerton, C. T., Hugenholtz, P. & Tyson, G. W. CheckM: assessing the quality of microbial genomes recovered from isolates, single cells, and metagenomes. (2014) doi:10.7287/peerj.preprints.554v1.

92. Sieber, C. M. K. et al. Recovery of genomes from metagenomes via a dereplication, aggregation and scoring strategy. Nat. Microbiol. 2018 37 **3**, 836–843 (2018).

93. Parks, D. H. et al. GTDB: an ongoing census of bacterial and archaeal diversity through a phylogenetically consistent, rank normalized and complete genome-based taxonomy. Nucleic Acids Res. 50, D785–D794 (2022).

94. Richter, M. & Rosselló-Móra, R. Shifting the genomic gold standard for the prokaryotic species definition. Proc. Natl. Acad. Sci. U. S. A. 106, 19126–31 (2009).

95. Arkin, A. P. et al. KBase: The United States Department of Energy Systems Biology Knowledgebase. Nat. Biotechnol. 2018 367 **36**, 566–569 (2018).

96. Langmead, B. et al. 2C-Ultrafast and memory-efficient alignment of short DNA sequences to the human genome. Genome Biol. 10, R25 (2009).

